# Directing multicellular organization by varying the aspect ratio of soft hydrogel microwells

**DOI:** 10.1101/2021.09.17.460849

**Authors:** Gayatri J. Pahapale, Jiaxiang Tao, Milos Nikolic, Sammy Gao, Giuliano Scarcelli, Sean Sun, Lewis H. Romer, David H. Gracias

**Affiliations:** Department of Chemical and Biomolecular Engineering, Johns Hopkins University, Baltimore, MD 21218, USA; Department of Mechanical Engineering, Johns Hopkins University, Baltimore, MD 21218, USA; Maryland Biophysics Program, Institute for Physical Science and Technology, University of Maryland, College Park, MD, 20742, USA; Fischell Department of Bioengineering, University of Maryland, College Park, MD 20742, USA; Institute of NanoBioTechnology (INBT), Johns Hopkins University, Baltimore, MD 21218, USA; Department of Cell Biology, Johns Hopkins School of Medicine, Baltimore, MD 21205, USA; Department of Anesthesiology and Critical Care Medicine, Johns Hopkins School of Medicine, Baltimore, MD 21205, USA; Department of Biomedical Engineering, Johns Hopkins School of Medicine, Baltimore, MD 21205, USA; Department of Pediatrics, Johns Hopkins School of Medicine, Baltimore, MD 21205, USA; Department of Center for Cell Dynamics, Johns Hopkins School of Medicine, Baltimore, MD 21205, USA; Department of Oncology, Johns Hopkins School of Medicine, Baltimore, MD 21205, USA; Sidney Kimmel Comprehensive Cancer Center, Johns Hopkins School of Medicine, Baltimore, MD 21205, USA; Department of Materials Science and Engineering, Johns Hopkins University, Baltimore, MD 21218, USA; Department of Chemistry, Johns Hopkins University, Baltimore, MD 21218, USA; Laboratory for Computational Sensing and Robotics (LCSR), Johns Hopkins University, Baltimore, MD 21218, USA

**Keywords:** tissue engineering, tubulogenesis, hydrogels, self-assembly, curved geometry, protein patterning

## Abstract

Multicellular organization with precise spatial definition is an essential step in a wide range of biological processes, including morphogenesis, development, and healing. Gradients and patterns of chemoattractants are well-described guides of multicellular organization, but the influences of three-dimensional geometry of soft hydrogels on multicellular organization are less well defined. Here, we report the discovery of a new mode of self-organization of endothelial cells in ring-like patterns on the perimeters of hydrogel microwells that is independent of protein or chemical patterning and is driven only by geometry and substrate stiffness. We observe quantitatively striking influences of both the microwell aspect ratio (ε = perimeter/depth) and the hydrogel modulus. We systematically investigate the physical factors of cells and substrates that drive this multicellular behavior and present a mathematical model that explains the multicellular organization based upon balancing extracellular and cytoskeletal forces. These forces are determined in part by substrate stiffness, geometry, and cell density. The force balance model predicts the direction and distance of translational cell migration based on the dynamic interaction between tangential cytoskeletal tension and cell-cell and cell-substrate adhesion. We further show that the experimental observations can be leveraged to drive customized multicellular self-organization. Our observation of this multicellular behavior demonstrates the importance of the combinatorial effects of geometry and stiffness in complex biological processes. It also provides a new methodology for direction of cell organization that may facilitate the engineering of bionics and integrated model organoid systems.

## Introduction

Multicellular spatial organization drives morphogenesis, development, healing, and homeostasis (*1–5*), and its disruption has been implicated in the onset of diseases and the development of pathobiological processes as diverse as hypertension and malignancy (*6–10*). Multicellular organization is regulated by various environmental cues, including biochemicals such as signaling molecules and transcription or growth factors (*11–14*) and physical parameters like shear, stiffness, and geometry (*15–20*). Researchers have reported that multicellular organization of vascular cells is controlled by the concentration and activities of biochemicals such as semaphorin 3E, platelet-derived growth factor (PDGF), and vascular endothelial growth factor (VEGF), as well as physical factors including mechanical stretch and shear. Alteration of some of these factors may lead to pathologies like endothelial hyperplasia and abnormalities in capillary shape, diameter, and permeability (*18, 21-24*). Similarly, developmental processes such as the formation of intestinal villi involves the sequential multicellular layered organization of intestinal cells in response to spatial gradients of differentiation and growth factors, leading to differential strain between layers and resulting in eventual folding that forms the villi (*25, 26*).

Recent advances in hydrogel synthesis and microscale patterning have led to discoveries of the roles of physical cues such as stiffness and geometry in cell organization (*27*). Studies have shown that only soft hydrogels with a stiffness less than 2 kPa support the spontaneous reorganization of epithelial cell aggregates to form lumenized structures like ducts (*8, 28*). Similarly, hydrogels with stiffness less than 1 kPa support endothelial tubulogenesis (*29–31*) and the formation of tumor spheroids, in contrast to the monolayers that form on stiffer substrates (*7*). The role of geometry in cell organization is most commonly probed by spatial confinement of cells to various adhesive (e.g., collagen, fibronectin) and non-adhesive (e.g., Pluronics) two-dimensional patterns (*32–36*). Recent studies have used three-dimensional (3D) geometries to study cell alignment and migration, but these have been done predominantly on substrates such as polydimethylsiloxane (PDMS) with non-physiological stiffnesses of >1 MPa (*37–39*). Hence the influence of geometry on multicellular organization in soft anatomically relevant 3D shapes is unclear. Here, we report a novel type of endothelial cell self-organization in the absence of any protein patterning, which occurs only within microwells of soft hydrogels with moduli in the range of 1-4 kPa. Rather than a spatial pattern of adhesive and less adhesive regions dictating substrate affinity, the self-organization we discovered is the consequence of the variation of two physical cues - namely stiffness and geometry. We observe that more than 90% of the cells on soft 2 kPa hydrogels migrate towards and organize on the edge of the microwells, whereas cells on stiffer 35 kPa hydrogels uniformly spread throughout the microwells (Fig. 1A). These findings contrast with studies reported with fibroblasts, stem cells, and epithelial cells, in which cells on stiff substrates avoided microwells(*37, 39*). We quantitatively characterized multicellular self-organization as a cell distribution ratio (DR) that is defined as the ratio of the number of cells on microwell edge (as the numerator) to the number of cells in the center (as the denominator). We report a strong correlation between the organization of cell populations and the microwell aspect ratio (ε), defined as the ratio of microwell perimeter to the depth on soft hydrogels. We investigate this phenomenon by measuring the DR in the context of different levels of cell contractility (cell tension), cell density, and cell adhesion and test our findings by implementing a force balance-based mathematical model. Apart from affecting multicellular organization in periodic microwells, we further show that geometric adjustments in soft hydrogels can be used to self-organize cells in a variety of CAD-designed microwell shapes, such as the letters of the word “CELL”. We believe that our discovery substantially advances the understanding of the complex role of geometry and stiffness on multicellular organization and will guide the design of microenvironments that direct multicellular organization for complex devices, mimetic tissues, and organoid designs in the absence of protein patterning.

**Fig.1.**
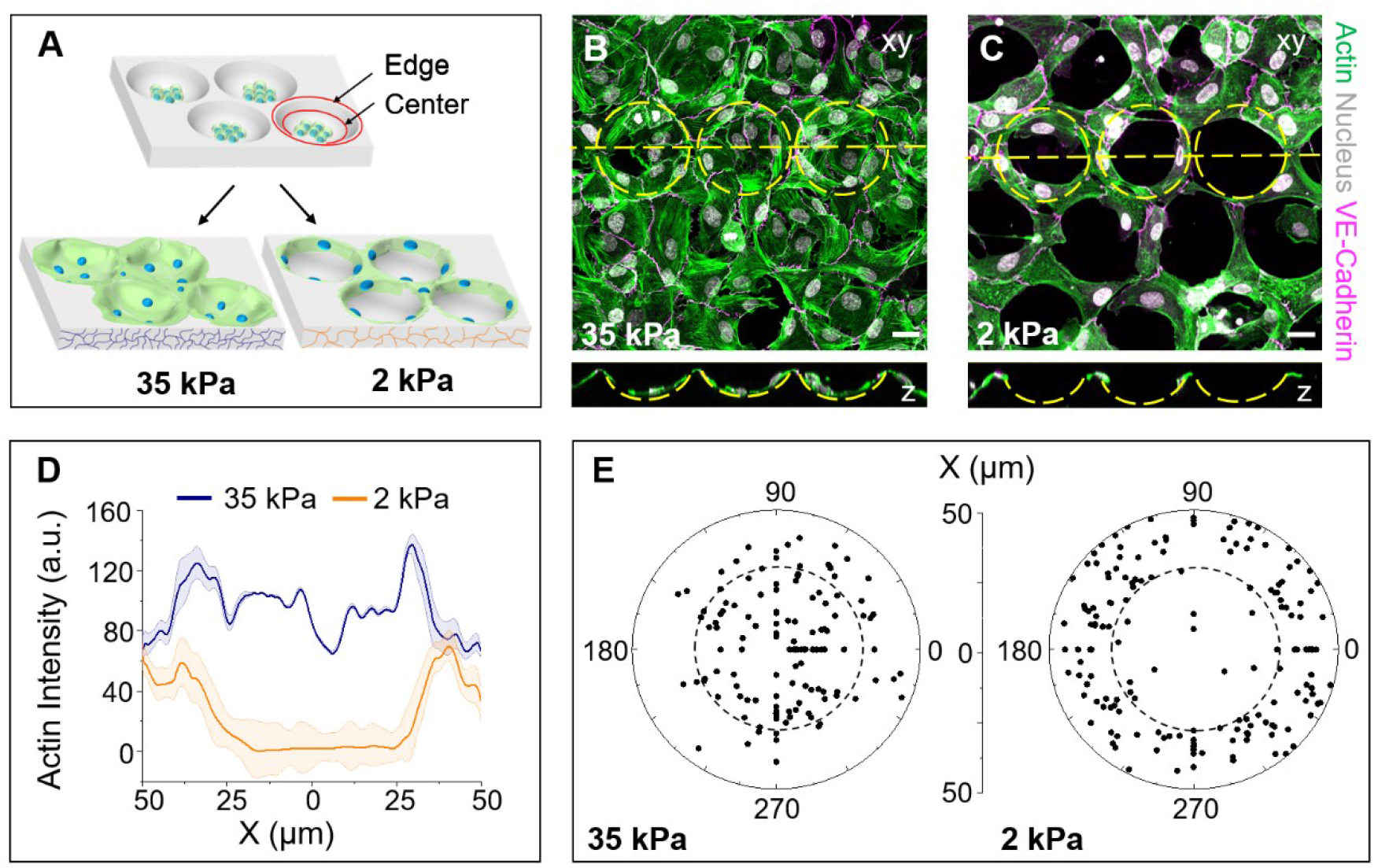
Multicellular organization is guided by geometry on soft 2 kPa hydrogels. (**A**) Schematic depicting our findings showing multicellular organization driven by geometry and stiffness in the absence of protein substrate patterns. We define two regions of equal projected area; edge (between the red circles) and center (inside the smaller red circle) to characterize the multicellular organization. (**B** and **C**) Confocal imaging showing the top (xy) and side-views (z) of actin (green), nucleus (gray) and VE-cadherin (magenta) stained cells 24 h after seeding in hemispherical microwells. The images show uniform multicellular distribution all over the stiff 35 kPa hydrogel microwell area (B). In contrast, we observe a distinct ring-like multicellular self-organization around the edge of the soft 2 kPa hydrogel microwell (C). Yellow dashed lines indicate the positions of the microwells and the axis along which the side view is shown. (**D**) Comparison plot of actin intensity measured along the microwell diameter (X), in stiff 35 kPa and soft 2 kPa microwells. The plot depicts the average intensity ± SEM for 100 microwells over three independent experiments. (**E**) Polar plot showing distribution of cells in the microwells quantified by measuring the distance of the nucleus from the center of the microwell. Region inside the dotted circle indicates the center region of the microwell. This plot was generated from measurements using 200 cells from four independent experiments. Scale bar = 25 μm.

## Results

### Endothelial cells self-organize into ring-like patterns on soft hydrogel microwells in the absence of protein patterns

We used photolithography and molding to pattern hydrogels of Young’s moduli 2 kPa and 35 kPa with microwells 80 μm in diameter and 20 μm depth. We chose these dimensions to mimic geometries found in anatomical structures, including arteries, mammary acini, intestinal crypts, alveoli, and epidermis (*40*). We arranged the microwells periodically in a hexagonal array where the spacing between two adjacent microwells is greater than or equal to 5 μm. We seeded human umbilical vein endothelial cells (HUVEC) in hydrogel microwells. We observed a striking difference in cell organization based on hydrogel stiffness: cells in stiff 35 kPa microwells organized into a uniform monolayer in and around the microwells (Fig. 1B), whereas cells in soft 2 kPa microwells self-organized exclusively (>90%) on the periphery of microwells in the shape of rings (Fig. 1C). We used gelatin hydrogels for the majority of our experiments and verified this self-organization with polyacrylamide hydrogels (Fig. S1).

Using live-cell imaging, we observed that HUVEC in soft 2 kPa microwells always started at the center of the microwells and then gradually moved to the edges of the microwell After 8 to 10 h of seeding, most of the cells in soft 2 kPa microwells were on the microwell periphery. While we observed that cells continued to move along the microwell edge for the remaining observation duration (cells are observed for a total of 24 h), we did not observe significant changes in the cellular organization beyond this time point and refer to it as the steady state.

We categorized the cells in the microwells into two regions of equal projected area: microwell edge and microwell center (Fig. 1A). We quantified the DR by counting the cells based on the nuclear position from the microwell center (detailed in the methods section). We observed that the steady-state DR for cells in soft 2 kPa microwells (DR=15) was approximately 30 times that for cells in stiff 35 kPa microwells (DR=0.5) (Fig.1E). In addition to DR, we also measured the distribution of average actin fluorescence intensity in the microwells by using phalloidin-labeled actin, which shows minimal (0±10 a.u. (arbitrary units)) actin in the central region with most (~70±10 a.u.) of the actin accumulating on the edges of soft 2 kPa microwells (Fig. 1D). In contrast, for stiff 35 kPa microwells, the actin signal is higher and observed all over the microwell (between 80-120 a.u.). The drop in intensity observed at the microwell center results from the nuclear clustering at the microwell center.

### The topography and mechanical properties of the hydrogel microwells are uniform

We characterized the topography and mechanical homogeneity of the gelatin microwells using scanning electron microscopy (SEM), optical microscopy, and Brillouin microscopy. SEM images taken at a tilt angle of 60° show that the surfaces of the soft 2 kPa and stiff 35 kPa microwells are uniformly flat and that there is no topographical irregularity (Fig. 2A, B). We also observed that the self-organization in soft 2 kPa microwells occurred consistently as verified over large microwell patterned areas (1 mm x 1 mm) using fluorescence microscopy, thus confirming that the microwell patterns are spatially uniform (Fig. S2).

**Fig. 2.**
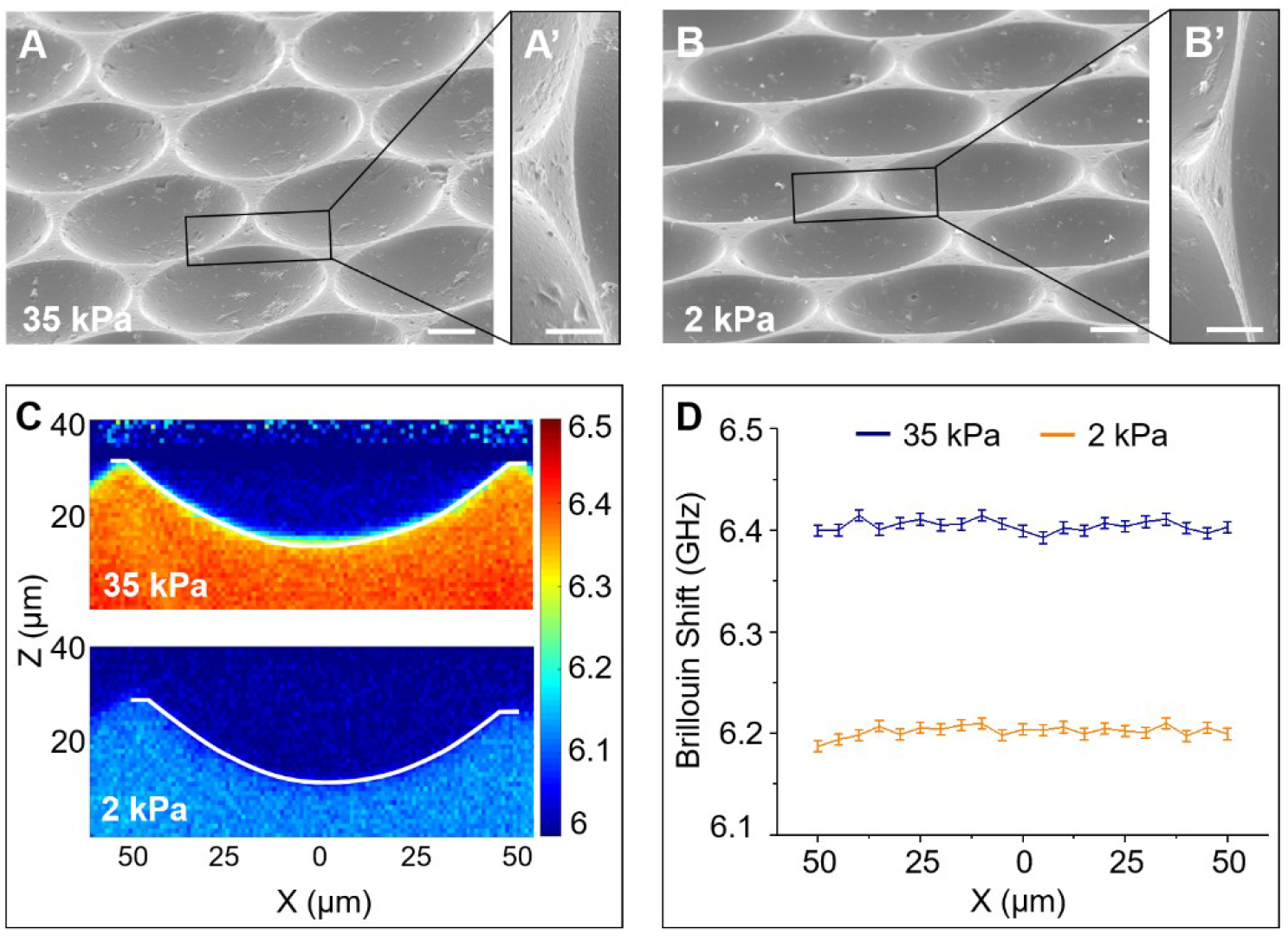
Microwells are topographically and mechanically homogeneous. (**A** and **B**) Tilted SEM micrographs of stiff 35 kPa and soft 2 kPa microwells with magnified sections (**A’** and **B’**) to focus on the edge of the microwells depicting that the surface topography of the microwells is homogeneous and similar around the edges. Scale bar = 10 μm for (A, B) and 5 μm for (A’, B’). (**C**) Representative Brillouin images of stiff 35 kPa and soft 2 kPa microwell with the white curved line indicating the boundary between PBS and the hydrogel. (**D**) Brillouin shifts on the microwell surface along the diameter (x) just below the hydrogel-liquid interface. Data represented is the mean ± standard deviation.

To ensure that mechanical heterogeneities were not driving cell migration to the microwell edge via durotaxis(*41, 42*), we mapped the local longitudinal moduli of the hydrogels using Brillouin microscopy(*43, 44*). The Brillouin frequency shift is sensitive to changes in material stiffness and has previously been used to characterize several types of soft materials, including gelatin-based hydrogels(*45*). We quantified the Brillouin frequency shift of the hydrogel cross-sections and found it constant within 0.08% relative error on nearly all microwell surfaces (Fig. 2C, D, and Fig. S3). These measurements confirm the homogeneous mechanical properties of the hydrogel.

### Self-organization occurs in soft 2 kPa microwells of different shapes

We studied multicellular self-organization in hemispherical, cylindrical, cubic, and pyramidal microwells. In all cases, we observed self-organization at the edges of only the soft 2 kPa microwells without significant bias in DR for any of the studied shapes (Fig. S4). This observation indicated that the circular shape of the microwell did not specifically induce the observed multicellular organization. Also, the fact that cells self-organized on the top edge of both hemispheres (one circular edge on the tops of the microwells) and cylinders (two circular edges, one at the top and one at the bottom of each microwell) indicated that contact guidance is not driving the observed self-organization (*46*). Following these observations, we used hemispherical microwells for all our further studies.

### Self-organization depends on the microwell aspect ratio

In our preliminary experiments, we observed that HUVEC formed a monolayer on flat soft 2 kPa hydrogels. This observation indicated that microwell depth is a driving factor in the self-organization of cells into ring-like structures. We, therefore, investigated the influence of microwell geometry on cell organization in detail by varying the microwell aspect ratio (*ε*) of soft 2 kPa microwells. We systematically varied the microwell depth from 25 μm to 0.5 μm while keeping the perimeter constant at 250 μm.

We observed that for microwells deeper than 10 μm (*ε* < 25), cells migrated away from the center to self-organize around the edge with a high DR that was greater than five (Fig. 3A, Fig. 3B red box, and Fig. S5A). For microwells less than 10 μm in depth (*ε* > 25), more cells tend to stay in the center region. The distribution of cells between the microwell edge and center approaches equality (DR→1) for microwells with depth less than 2 μm (Fig. 3A, Fig. 3B yellow box, and Fig. S5B). A similar trend was observed for soft 2 kPa microwells with 470 μm perimeter (Fig. S6). These observations indicated that the microwell aspect ratio is a critical factor in determining cell organization.

**Fig. 3.**
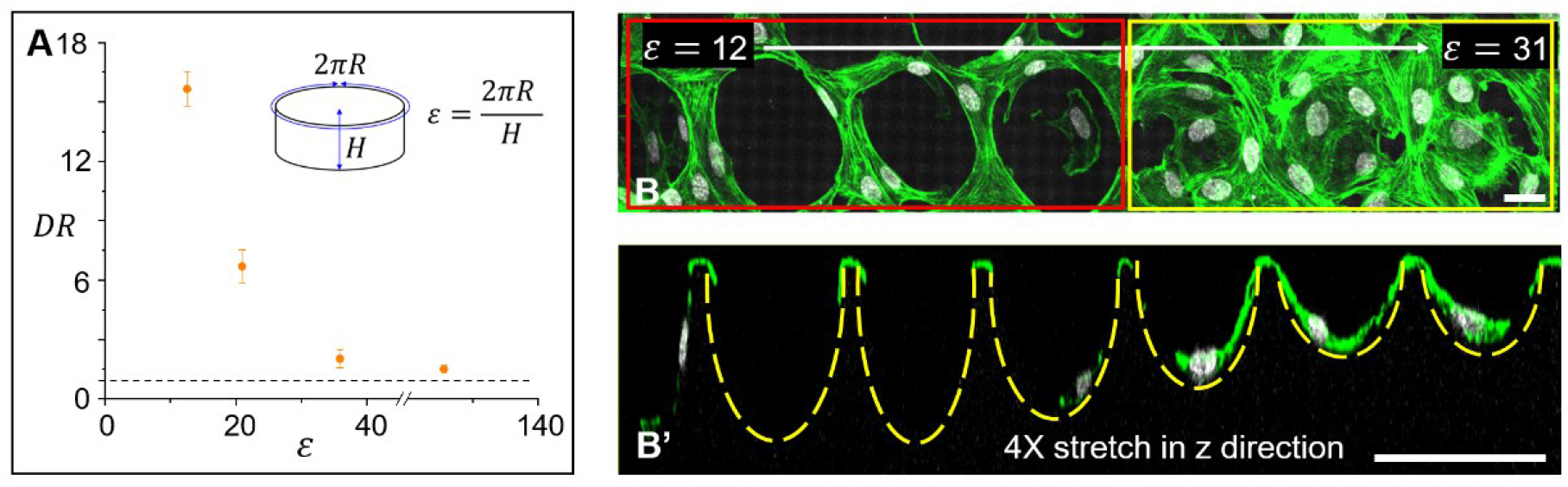
Multicellular organization in 2 kPa microwells depends on microwell aspect ratio. (**A**) Plot depicting the relation between multicellular organization as quantified by DR and the aspect ratio ε, for the soft 2 kPa microwell geometry. Data presented are quantified from 100 cells for each aspect ratio and repeated three times. Error bars indicate standard deviation between each experiment. (**B** and **B’**) Confocal imaging showing the top (xy) (B) and side-views (z) (B’) of cells stained for actin (green) and nucleus (gray) in soft 2 kPa microwells with decreasing depth (increasing ε) from left to right. Red box depicts microwells with depth > 10 μm (ε < 25) and yellow box depicts microwells with depth < 10 μm (ε > 25). The image in panel (B’) is stretched by a factor of four in the z-direction, to clearly visualize the shallower microwells. Yellow dashed arcs are included to depict the microwell shape in the z plane. Scale bar = 25 μm.

### Multicellular self-organization occurs at an optimal cell density

We observed that the cell movement toward the edges of soft 2 kPa microwells strongly depended upon the cell seeding density. At high seeding densities (>5 × 10^4^ cells/cm^2^), cells occupied the entire microwell area in both stiff 35 kPa and soft 2 kPa microwells and did not organize in ring-like structures. However, we found that the cells kept moving in the central region of the stiff 35 kPa and the soft 2 kPa microwells when they were seeded at lower densities (~6.5 ×10^3^ cells/cm^2^) (Fig. S7A,B). Interestingly, when independently moving and noncontiguous cells in soft 2 kPa microwells encountered one another, they tended to move toward the microwell edge. For cells in soft 2 kPa microwells at a very low density, the DR was less than one and decreased as the microwell depth increased (or as ε decreased), suggesting that cell migration toward the microwell edge may be enhanced by the presence of neighboring cells (Fig. S8). We found that the optimal seeding density that allowed the cells to self-organize on the soft 2 kPa microwell edge was between 2-3 ×10^4^ cells/cm^2^. At this optimal density, cells seeded in soft 2 kPa microwells moved toward and along the edges and exhibited greater translational motion than cells in stiff 35 kPa microwells (Fig. S7C and D). These observations suggested that cell-cell interactions play a role in cell self-organization at the microwell edges in 2 kPa microwells.

### Delineating the role of substrate stiffness on multicellular self-organization

Multicellular self-organization into ring-like structures only occurred in soft 2 kPa microwells and not in stiff 35 kPa microwells. This observation indicates that substrate stiffness is a regulating factor in guiding the observed self-organization. It is known that substrate stiffness affects cell size, cytoskeletal contractility, and cell adhesions (*47,49*). Since stiffer substrates lead to a larger spreading area which may lead to the filling of the microwells, we first investigated the cell size in the microwells. We measured the cell area by projecting the confocal z-stack on a plane and found that the sizes of the cells in 35 kPa and that of the cells in 2 kPa microwells were similar (Fig. S9). We thus hypothesized that this difference in behavior could in part be due to the difference in cytoskeletal contractility on the two hydrogels. Previous studies have established that substrate stiffness can be correlated to cell contractility and that cell contractility increases as substrate stiffness increases (*47–49*). We tested this hypothesis by altering the cell contractility using pharmacological agents.

Specifically, we suppressed Rho-activated, myosin II-dependent contractility of the microfilament cytoskeleton in cells seeded in stiff 35 kPa microwells by inhibiting ROCK with Y27632 (30 μM). In cells seeded in soft 2 kPa microwells, we inhibited myosin dephosphorylation and inactivation using the phosphatase inhibitor Calyculin A (0.1 nM) to increase the contractile state of the microfilament cytoskeleton. We observed that ROCK inhibition in cells seeded to stiff 35 kPa microwells led to self-organization on the edge of microwells similar to that seen in untreated cells on soft 2 kPa microwells (Fig. S10A). Interestingly, reciprocal conditions whereby Calyculin A treated cells were seeded in soft 2 kPa microwells yielded the accumulation of these cells in the microwell centers in a manner that recapitulated the behavior of untreated cells on stiff 35 kPa microwells (Fig. S10B). These results support the hypothesis that cytoskeletal contractility (as dictated by the substrate stiffness) is a significant contributor in directing cell self-organization and that stiffer substrates inhibit the geometry-enhanced cell motility observed in microwells comprised of softer substrate. Next, we investigated the effect of substrate stiffness on cell-substrate adhesion as described in the following section.

### Influence of cell-cell and cell-substrate interaction on multicellular self-organization

Cell-cell and cell-substrate interactions are strongly influenced by stiffness and cell density. Prior reports show that increased substrate stiffness promotes robust cell-substrate interaction through focal adhesions (FA) due to force loading and increased protein clustering (*50, 51*), whereas increased substrate stiffness destabilizes VE-cadherin mediated cell-cell junctions and weakens cell-cell interactions (*52, 53*). We quantitatively investigated the effect of the regulating parameters, microwell stiffness, size, and cell density on cellular interactions and their impact on our observed multicellular self-organization using immunostaining.

We began by studying paxillin in focal adhesions at 3 h and 24 h after cell seeding to quantify the 3D FA surface area and volume. We grouped the FA into two categories - microwell edge or microwell center. FA produced by cells that extended from the microwell edge to the microwell center were categorized in the appropriate group based upon the FA location. We also studied VE-cadherin at EC cell-cell junctions after 24 h of seeding in soft 2 kPa and stiff 35 kPa microwells.

We observed that cells in stiff 35 kPa microwells established larger, streak-like FA. At 3 h, FA produced at the edges of the 35 kPa microwells were larger than those in the microwell center (Fig. S11A-i), and this difference resolved by the end of 24 h (Fig. 4A-i, c). In contrast, at both 3 h and 24 h time points FA in HUVEC at the center of the soft 2 kPa microwells were smaller and predominantly punctate structures (Fig. S11B-i and Fig. 4B-i, red box), while those at the edge were streak-like FA and ~ 2.5 times larger than those in the center (Fig. S11B-i and Fig. 4B-i, yellow box). Moreover, FA size on the edges of the soft 2 kPa microwells was comparable to the FA size measured in stiff 35 kPa microwells at 24 h (Fig. 4C and Fig. S12A, B). A similar distribution of FA was observed in cells seeded at low and very high density in the soft 2 kPa microwells (Fig. S12C, D-i).

**Fig. 4.**
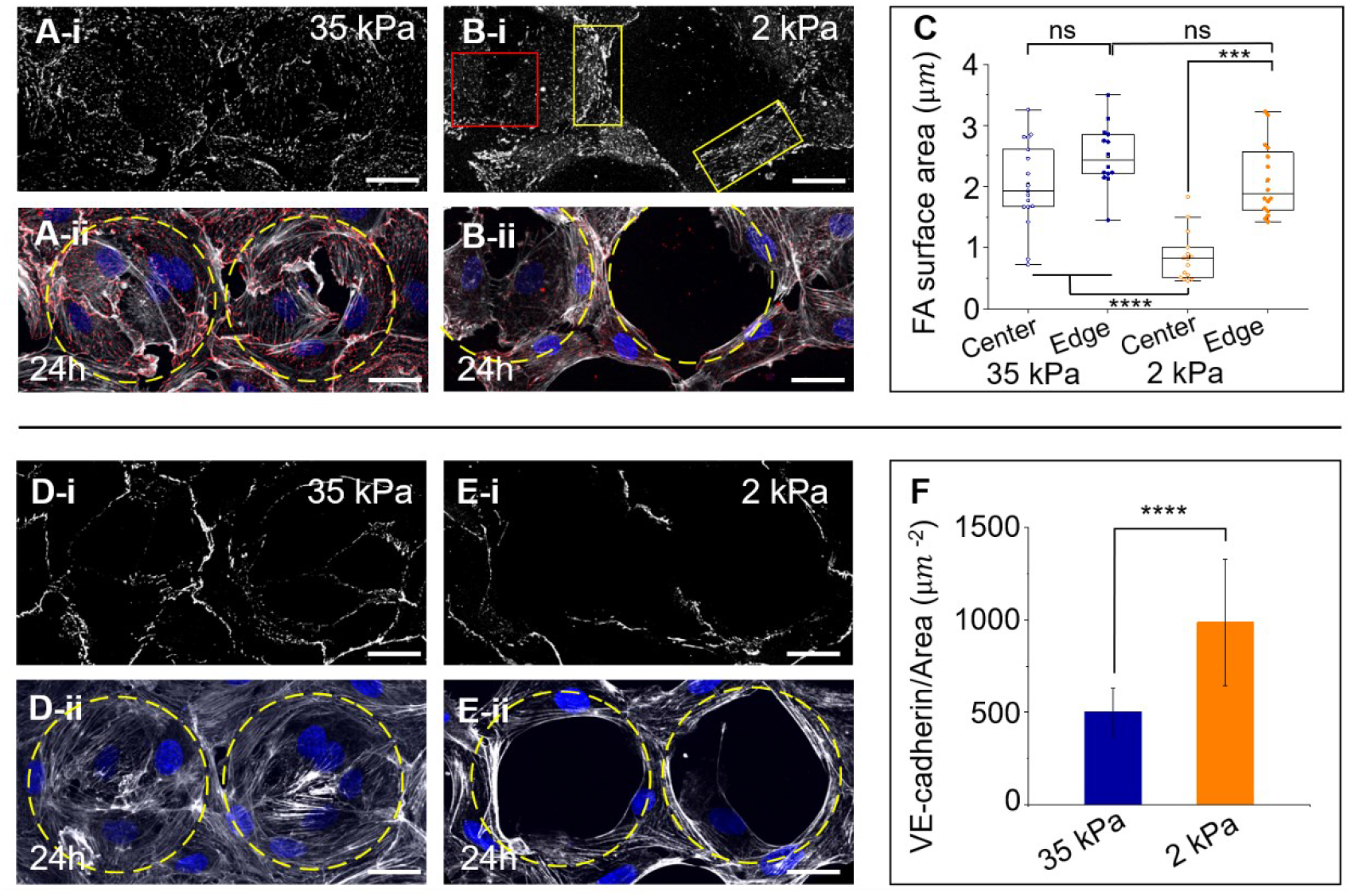
Cell-substrate and cell-cell interactions at the center and edge of soft and stiff microwells. (**A** and **B**) Confocal imaging showing the top view of cells immunolabeled for paxillin (gray) 24 h after seeding in stiff 35 kPa and soft 2 kPa microwells respectively (A-i, B-i) and the corresponding composite top view of cells labeled for paxillin (red), actin (gray), and nucleus (blue) (A-ii, B-ii). The yellow box shows the streak-like FA on the edge and the red box shows the dot-like FA inside the soft 2 kPa microwell. (**C**) Box plot depicting the FA size for cells at the center and edge of the stiff 35 kPa and soft 2 kPa microwells. Data presented are for n ≥ 15 cells pooled from three independent experiments. ****denotes *P* values < 0.0001 and *** denotes *P* value < 0.001 calculated by one-way ANOVA followed by Tukey’s multiple comparison. (**D** and **E**) Confocal imaging showing the top view of cells labeled for VE-cadherin (gray) 24 h after cell seeding in stiff 35 kPa and soft 2 kPa microwells respectively (D-i, E-i) and the corresponding top view for cell stained for actin (gray) and nucleus (blue) (D-ii, E-ii). (**F**) Plot depicting average VE-cadherin intensity per unit junction area. Data presented are for n>10 sample areas pooled from three independent experiments. **** denotes *P* value < 0.0001 calculated by Student’s *t*-test. Yellow dashed circles indicate the microwell position. Scale bar = 25 μm.

Thus, the limited translational motion of the cells in the 35 kPa microwells may be attributed in part to the presence of large FA made by cells inside microwells (Fig. S7D). In contrast, cells formed much smaller FA inside softer (2 kPa) microwells. This facilitated their motility out of the microwell center and toward the edges, where they established large FAs and stabilize their position (Fig.S7C). Since inhibition of ROCK reduces the formation of FA due to reduced stress fiber formation (*54*), Y27632 caused cells in 35 kPa microwells to mimic the behavior of cells in 2 kPa microwell centers and migrate to the edge.

VE-cadherin localized along the majority of each cell’s perimeter in stiff 35 kPa microwells, as HUVEC there are tightly contiguous to neighboring cells. In contrast, cells in soft 2 kPa microwells only exhibited cell-cell contacts along ~40-50% of their perimeter (Fig. 4D, E). We quantified the VE-cadherin intensity and normalized these values per unit of junctional area and were surprised to observe that the normalized VE-cadherin intensity for cells in soft 2 kPa microwells was approximately twice that seen in cells in stiff 35 kPa microwells (Fig. 4F). Thus, in addition to forming larger FA, cells at the edges of the soft 2 kPa microwells also formed more cell-cell junctions that were heavily populated by VE-cadherin.

Thus, it is evident from our experiments that the multicellular self-organization that we observe involves an interplay between cell-cell and cell-substrate interactions and that this dynamic is influenced by the three parameters noted above: substrate stiffness, microwell aspect ratio, and cell plating density.

### Balance between cell tension, adhesion, and cohesion can explain the observed cell organization

We developed a mathematical model to integrate our observations and gain insight into our observed multicellular self-organization (see Supplementary text for full description). We modeled force-balance relationships and active force generation by cells to simulate the dynamics of two cells moving along the microwell wall with a depth *H* and radius *R* (Fig. 5A and Fig. S13A). We assume that the all the forces act only on each cell’s endpoint (nodes). Thus, a line segments can be used to simplify the cell shape and give a good approximation of force condition on the cell. Complicated assumptions for cell shape, such as a curved line or a 3D shape may not be necessary as these models would yield similar results because the forces are concentrated on cell endpoints. We can write the force-balance equation at *i* ^th^ endpoint of *j* ^th^ cell, *x_j,i_*, as:

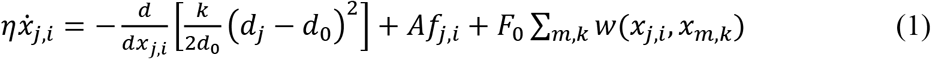

where the left-hand of the equation is the cell-substrate interaction (adhesion) described as the sliding frictional force (*F_η_*) between the cell and the substrate with coefficient *η*. The right-hand side describes cell active forces comprising volume (length) regulation by cellular cytoskeletal forces (*F_k_*) and active protrusion/contraction. The last term on the right-hand side of equation (1) describes the cell-cell interaction (cohesion) (*F_w_*).

**Fig. 5.**
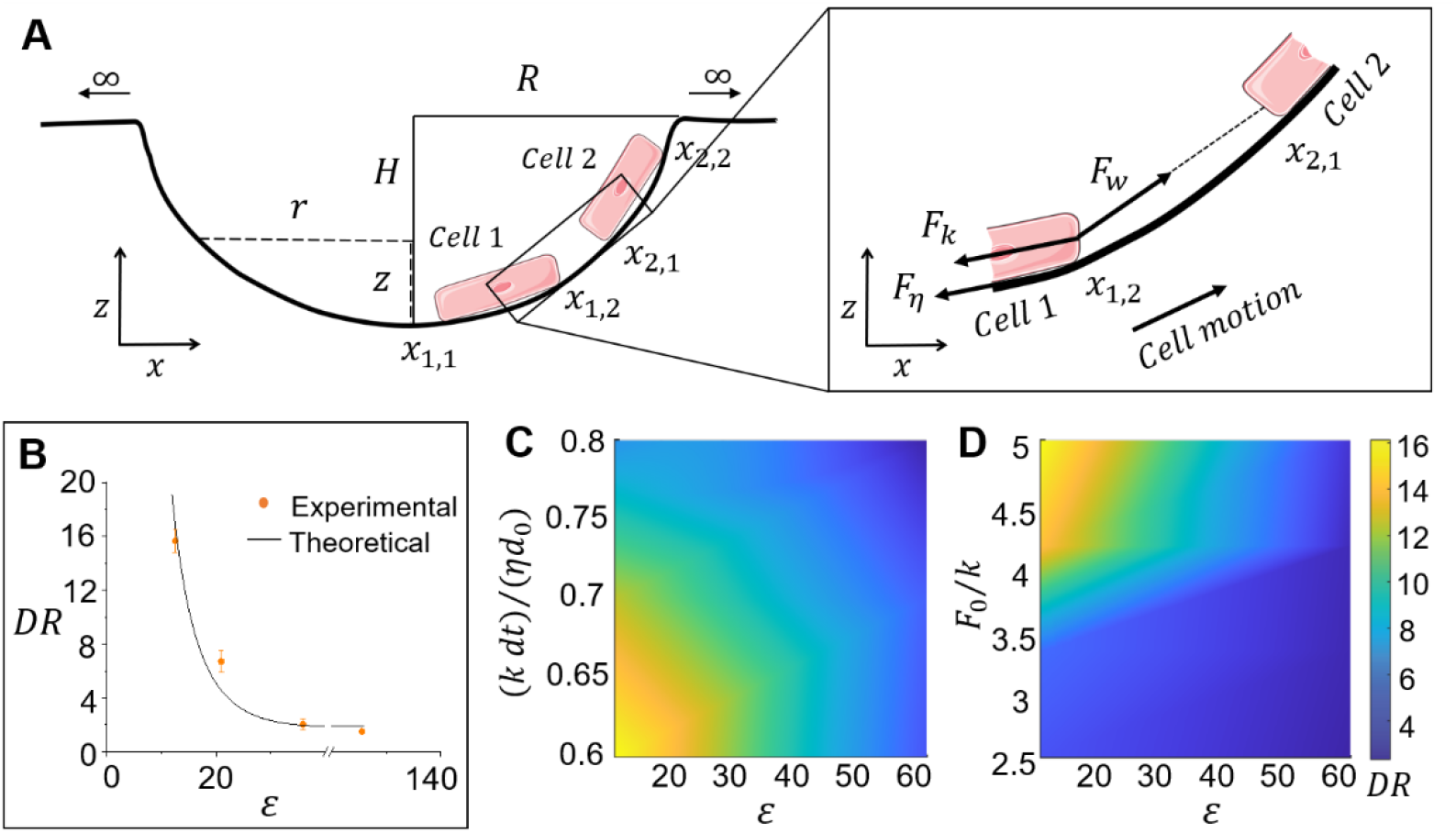
Force balance model predicts the observed pattern of multicellular organization. (**A**) Schematic depicting the force balance model parameters. The model analyses the cell tension (*F_k_*), cohesion (*F_w_*), and adhesion (*F_η_*) forces, assumed to be concentrated at the end nodes of the cells, to predict the observed multicellular organization. (**B**) Plot depicting the experimental observations and theoretical predictions for the relationship between cell arrangement as characterized by DR and microwell aspect ratio. (**C** and **D**) Plot showing the DR for various aspect ratios as a function of cytoskeletal tension (C) and cell-cell interaction (cell density) (D).

We describe the cell cytoskeletal tension mechanics using a spring potential with the stiffness of *k*, which is directly related to the cells’ cytoskeletal stiffness, equilibrium length (*d*_0_), and current length (*d_j_*). The active protrusions and contractions, *f_j,i_*, generated by each cell is modeled by a random Gaussian noise term with zero mean, and scaled by a constant *A*. We assumed that the interaction force between the *i* ^th^ node of the *j* ^th^ cell and the *k* ^th^ node of the *m* ^th^ cell, *w*(*x_j,i_, x_m,k_*), mediated by VE-cadherin, follows a van der Waals-like potential that contains short-range repulsion and long-range attraction (Fig. S13B). The cell-cell interaction strength is related to cell seeding density, as well as the overall expression of VE-cadherin, and is described by a scaling constant *F*_0_ in our model. This interaction force is along the direction connecting the two cell endpoints: *x_m,k_* – *X_j,i_*.

This force equation of motion can explicitly compute the cell velocity and indicates that any unbalanced force between tension, cell-cell interaction, and random protrusions at each node must be compensated by the local motion along the microwell. Since the cell active forces are along the cellular orientation, the model suggests that the force is most likely to balance when cellular orientation is aligned with the local tangential direction. Specifically, the cell orientation is roughly along the local tangential direction towards the top edge in deeper microwells (*ε* < 25) and progressively shifts towards the center as the microwells become shallower (*ε* > 25); thus, the cells are more likely to stay near the top edge in deeper microwells while towards the center in shallower microwells (Fig. 5B) explaining the observed migration of cells to the edges of microwells.

Our model also explains the cellular organization on the microwell when cellular active forces, cell-cell, and cell-substrate interactions are varied to account for changes in microwell stiffness and cell density. Usually, higher cell stiffness, *k*, or lower cell-cell interaction strength, *F*_0_, requires a smaller tangential component of overall cellular tension for the force balance to be equal. Therefore, when cells are placed in stiff 35 kPa microwells, the resulting higher cytoskeletal stiffness and lower VE-cadherin activities than cells seeded in soft 2 kPa microwells tend to move the cells more towards the center to compensate for the extra increase in cell tension even in deeper microwells (Fig. 5C). On the other hand, when cells are seeded at a higher density in 2 kPa microwells, which increases the cell-cell interaction strength, *F*_0_, a higher tangential component of cell tension, is needed to balance the increased interaction, leading to cell migration towards the edge of the deeper microwell. Thus, when cells in soft microwells contact a second cell, the cells move more towards the top edge in microwells with a favorable aspect ratio and greater depth (Fig. 5D).

## Discussion

The physiological landscape features niches with heterogeneous mechanical and geometrical properties. These properties elicit diverse cell-extracellular matrix (ECM) interactions such as the cycling of cell adhesions and cytoskeletal stiffness changes that influence cell aggregation and tissue organization (*55, 56*). Our ability to create 3D micropatterns in hydrogels as soft as 2 kPa allows us to recreate anatomically relevant features that drive striking differences in self-organization using materials with physiologic stiffness. Our approach contrasts with most prior studies that utilize PDMS, which has a substantially higher Young’s modulus than materials that characterize most biological tissues (*38, 39, 57, 58*).

We have described a new mode of multicellular organization in response to physical cues that regulate cell migration and tissue morphogenesis. Specifically, we show that 3D geometry on soft hydrogels at cellular length scales can drastically influence the self-organization of endothelial cells (EC). We show that cells demonstrate a predilection to stay in locations where they can form stable cell-substrate adhesion and strong cell-cell interactions and explain it mathematically by considering the changes in the force balance between cytoskeletal tension and extracellular forces that include cell-cell and cell-substrate adhesion. This force balance can be adjusted by substrate stiffness, microenvironmental geometry, and cell density.

Studies have demonstrated that the physical properties of ECM and cell density both affect EC stiffness and EC interactions with substrate and with other cells. EC on stiffer matrices have higher cytoskeletal stiffness than on softer matrices (*48*). The individual EC is also stiffer than those in clusters, and decreasing cell-cell adhesion in groups of EC increases cell stiffness (*59*). Further, EC on stiff substrates make larger and stronger FA than those on soft substrates and do not undergo tubulogenesis (*29*). EC moving in pairs on stiffer substrates generate higher traction force (a measure of cell-substrate interaction) than on soft substrates, and traction forces generated by independently moving EC are lower than that generated by cell pairs in either environment (*60*).

Our discovery demonstrates that cells in 3D geometries on soft substrates integrate these responses to extracellular cues to self-organize in unique patterns. An intriguing finding in our study is that the geometry of the microwell - specifically the aspect ratio (*ε*), which is the ratio of the microwell perimeter and the microwell depth - impacts the magnitude of cytoskeletal tension that balances the extracellular interactions and hence multicellular organization. This impact of the microwell aspect ratio is due to the balance of cell-intrinsic and extracellular forces in the tangential direction at steady state (Fig. 5A). Cells at optimal density (2-3 ×10^4^ cells/cm^2^) in soft microwells have lower overall cytoskeletal tension, lower cell-substrate interaction at the centers than on the edges of the microwells, and higher cell-cell interaction. This imbalance drives the cell motion to the edge, where the larger tangential component of cell tension and higher traction (friction) generated due to cell-substrate interaction balance the cell-cell interaction. In contrast, cell-cell interactions in stiff microwells can be balanced at any location in the microwell due to the elevated levels of cell tension and cell-substrate interaction. We demonstrate that perturbing any of these three forces drastically changes the cell organization. Increasing cell tension in soft microwells by either increasing cell contractility or reducing cell density results in the loss of self-organization in soft microwells. The opposite is true for cells in stiff microwells, where decreasing cell tension leads to self-organization on the edges of the microwells (Fig. 5 and S13).

Such perturbations *in vivo* can lead to lethal pathologies. For example, EC organize into shorter or severely tortuous blood vessels in pathological conditions like hypertension, diabetes, and other vascular conditions with altered vascular wall mechanics (*61–63*). The inability of the cells to organize into healthy blood vessels is, in part, attributed to the stiffening of the ECM. Also, in the case of tumor blood vessels, elevated cell proliferation has been reported to affect cell organization resulting in randomly structured and leaky blood vessels that lack the hierarchical structure of a normal vasculature (*64*). Our experimental and theoretical insights into the mechanics that modulate morphogenesis may be harnessed to design *in vitro* microenvironments to study diverse cellular populations and their organization during normal development, aging, and disease pathogenesis.

From a technological perspective, our discovery’s key significance is that cell organization can be directed by defining the aspect ratio of the 3D geometry on soft hydrogels without the need for patterning cell adhesive and repellent regions. It is noteworthy that most prior literature utilizes patterns of chemoattractants or mechanical properties on flat substrates to manipulate cell organization (*34, 41*). In contrast, we guide multicellular organization without protein patterning. Thus, our approach to cell patterning offers a unique advantage to study cell organization in 3D and can provide avenues for co-culture systems to understand the interaction between endothelial cells and other cell types.

Our biofabrication approach utilizes photo and soft-lithography and is amenable to creating customizable shapes of hydrogel microwells. Thus, we can direct multicellular self-organization without protein patterning in various CAD-designed patterns. We highlight this characteristic by seeding cells on soft hydrogel microwells in the shape of the word “CELL” (Fig. 6 and Fig. S14). It is evident from the depth-coded 3D image stacks that the cells can self-organize even along these non-periodic and asymmetric micropatterns with good fidelity.

**Fig. 6.**
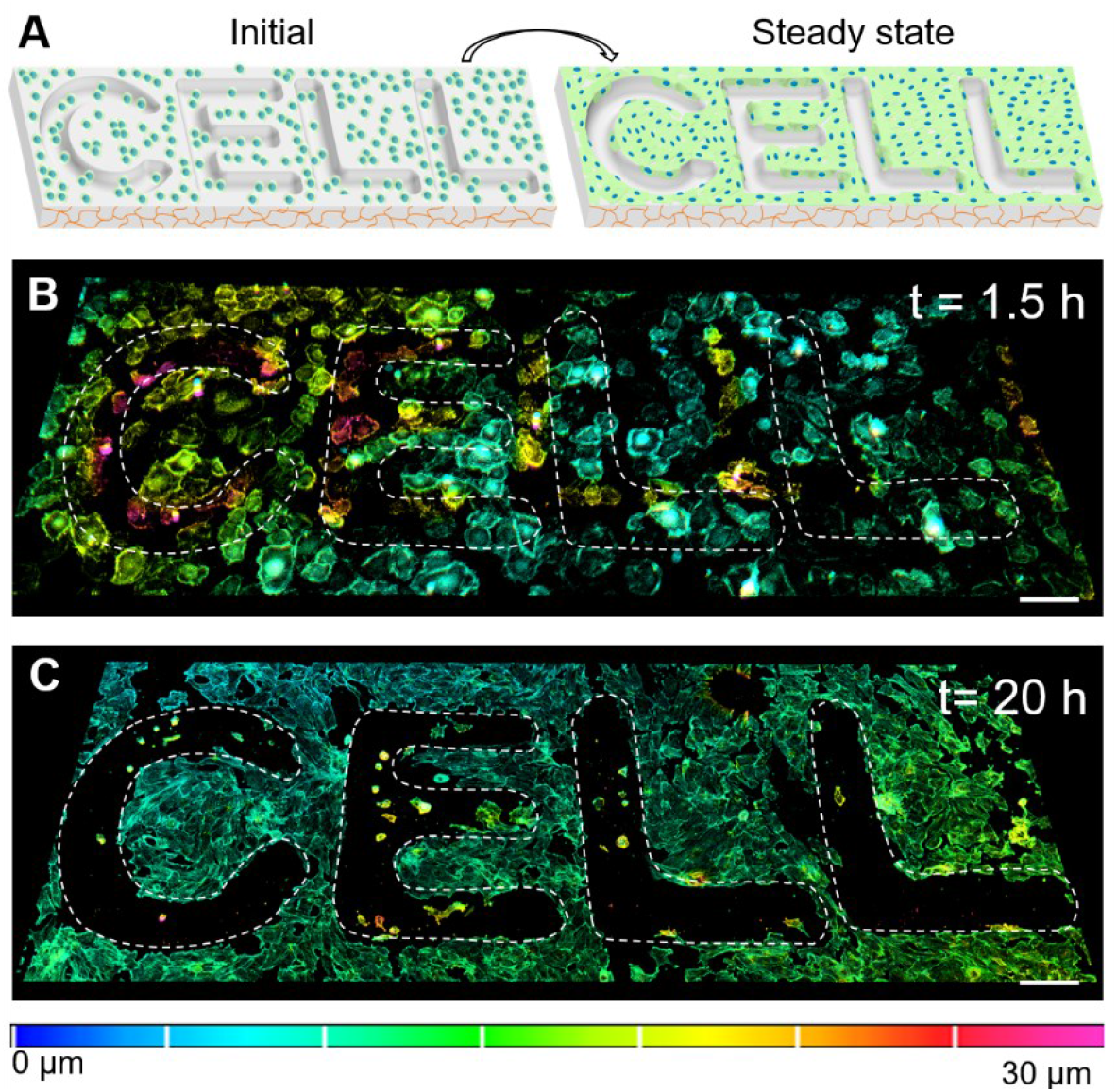
Directing multicellular organization in a predefined customizable pattern driven only by substrate geometry. (**A**) Schematic representing initial and steady state cell distribution on soft 2 kPa hydrogels patterned with the word CELL. (**B**) Depth coded confocal 3D stack of LifeAct-GFP-expressing HUVEC on soft 2 kPa hydrogel patterns 1.5 h (initial) after seeding. See Fig. C14B for the brightfield image which shows all the cells. (**C**) Depth coded confocal 3D stack of the same LifeAct-GFP-expressing HUVEC in (B) which were fixed and stained for actin to visualize all the cells on soft 2 kPa hydrogel patterns 20 h after seeding (t > steady state). Scale bar = 100 μm.

In conclusion, our findings provide a new paradigm to guide the patterning and self-organization of endothelial cells. We have defined the type and magnitude of the physical environmental cues required to drive self-organization. This may now be used to investigate fundamental questions in developmental biology, morphogenesis, and the etiology of diseases like cardiovascular conditions or cancer progression. In the future, patterns could also be utilized to self-organize blood vessel networks in the desired configurations by patterning the geometry of soft hydrogels without any superimposed surface chemical modifications. Since our approach allows highly parallel, reproducible, and cost-effective hydrogel patterning, we anticipate widespread applicability in cell and tissue engineering, cell signaling, and drug discovery. Finally, our discovery that striking differences in multicellular self-organization are caused by the combined effects of stiffness and microwell aspect ratio suggests that stiffness has a critical role in previously observed topography- and curvature-guided cell behaviors (*38, 39, 57, 65, 66*).

## Materials and Methods

### Fabrication of micropatterned hydrogel substrates

We fabricated the micropatterned hydrogels as reported previously (*40*). Briefly, we used appropriately thick positive tone photoresists (AZ9260 and AZ5214E, MicroChemicals GmbH and SPR220, Kayaku Advanced Materials) depending on the microwell depth and exposed them to UV light through a photomask to create micropatterns on a silicon wafer. After exposure and development, the micropatterns were heated beyond the photoresist glass transition temperature to induce photoresist reflow and obtain micropatterns with a curved profile. We used two steps of PDMS (Sylgard™ 184, Dow^®^) molding against the photoresist patterns to yield a final reusable micropatterned mold for gelatin. We sterilized the micropatterned molds under UV for 1 h and then treated them with 1% Bovine Serum Albumin (BSA) (Sigma-Aldrich) for 1 h at room temperature to prevent the hydrogel from sticking to the PDMS mold. We used gelatin (Type A from porcine skin, Bloom value 90-110 and 300, Sigma-Aldrich) to prepare the hydrogels. We prepared 12.5% wt/wt gelatin which was aseptically mixed with 1 mL of 10U/g-gelatin of microbial transglutaminase (Ajinomoto) to render the hydrogel thermostable. We waited for 9 h for the gels to crosslink and then stored them in PBS until used for the experiments.

### Cell culture, cell seeding, and live-cell imaging

We cultured Human Umbilical Vein Endothelial Cells (HUVEC) (ScienCell) in endothelial cell growth media (PromoCell) with 1% penicillin-streptomycin. We placed the hydrogels inside glass cloning rings (Fisher Scientific™) to seed the cells on the micropatterned hydrogels, which fit snugly around the hydrogels (*40*). We seeded the HUVEC at the desired cell density on the hydrogels and monitored their growth for 24 h. We treated the cells with the relevant pharmacological agent and its corresponding vehicle controls for experiments involving alteration of cytoskeletal stiffness. Y-27632 (Cell Signaling Technology) was included with the cells upon seeding, whereas Calyculin A (Sigma-Aldrich) was added 4 h following cell seeding.

We recorded the cell migration in the microwells at regular intervals for more than 18 h using a ×10/0.45 NA Ph1 objective on an inverted Nikon Eclipse Ti microscope (Nikon) with automated controls (NIS-Elements, Nikon) equipped with an incubation chamber. We generated the cell migration tracks using the MTrackJ plugin in ImageJ (NIH).

### Immunofluorescence experiments

We fixed the cells seeded in hydrogel microwells with a 3% paraformaldehyde (Sigma-Aldrich) for 20 min, permeabilized with 0.1% Triton X-100 (Sigma-Aldrich) for 15 min, and blocked with 2% BSA (Sigma-Aldrich) or 5% Normal Horse Serum (Invitrogen) in PBS for 1 h to prevent non-specific binding of the antibody used. After blocking, the samples were incubated for 1 h at 37°C with an appropriate primary antibody or fluorescent probe, followed by incubation at room temperature with a secondary antibody for 1 h. All antibodies and fluorescent probes were diluted in 0.1% BSA in PBS. Primary antibodies included rabbit anti-VE-cadherin antibody (160840, Cayman Chemicals,1:150) and rabbit anti-Phospho-Paxillin (69363, Cell Signaling, 1:100). The secondary antibody was goat anti-rabbit antibody conjugated with Alexa Fluor 488 (A-11034, Invitrogen™, 1:150 for VE-Cadherin and 1:100 for Paxillin). Actin was labeled with Alexa Fluor™ 488 Phalloidin (A12379, Invitrogen™), or Rhodamine Phalloidin (R415, Invitrogen™). Nuclei were labeled with 4’,6-Diamidino-2-Phenylindole, Dihydrochloride (DAPI) (D1306, Invitrogen™). We labeled the cells directly on the micropatterned hydrogels in the glass cloning rings to avoid disrupting the cell organization and stored them in PBS.

### Image acquisition and analysis

We detached the glass rings enclosing the hydrogel sample for imaging the cells in the microwells and placed the hydrogel samples either on glass-bottom dishes (MatTek Corporation) with the patterned surface close to the objective lens or mounted them in FluorSave (Millipore). We used a Nikon A1 confocal microscope to acquire the z-stacks and analyzed the stacks using ImageJ (NIH) and Imaris (Bitplane) software.

We used the confocal z-stack to quantify the DR by measuring the distance between the microwell center and the center of the nucleus. Cells overlapping both the central and edge regions were categorized to be located on the edge if >50% of the nucleus was present in the edge region and in the center if >50% of the nucleus was present in the center region. Nuclei with equal areas in both the regions were categorized based on the projected cell cytoskeleton area.

For measurement of paxillin-containing FA, we used the confocal z-stack to convert the paxillin signal into 3D surfaces while removing the paxillin signal that was not at the cell-substrate interface using the Imaris software. FA were categorized into two groups: microwell center or microwell edge. For VE-cadherin, we identified the junction region (or the region of interest, ROI) from the maximum intensity projection and used the same ROI with the average intensity projection to quantify the fluorescence intensity of VE-cadherin junctions using ImageJ. We verified these measurements in 3D using Imaris.

### Brillouin microscopy measurements

We measured the Brillouin shift of the hydrogels using a two-stage Virtually Imaged Phase Array (VIPA) spectrometer as described previously (*44, 67*). In brief, the sample was illuminated with 660 nm laser light through a 60x, 0.7NA objective (Olympus). Backscattered light was collected through the same objective and coupled through an optical fiber into a two-stage VIPA spectrometer. The optical spectra were recorded using an EMCCD camera (Andor). The sample was placed on a motorized stage on the Olympus IX81 microscope and scanned in 3D. One spectrum (50 ms exposure time) was recorded for each pixel (0.5 μm) in the confocal images. Data collection was performed through a custom LabView program. Each spectrum was fitted using the least squares method to localize Lorentzian-shaped Brillouin scattering peaks (MATLAB). The spectral dispersion (GHz/pixel) and free spectral range (FSR) were calibrated by measuring two materials of known Brillouin shift (water and methanol). The Brillouin shift’s spectral precision was typically 8 MHz (estimated as the standard deviation of the Brillouin shift of PBS shift in each image). Hydrogels were kept hydrated by immersion in PBS in a glass-bottom imaging dish (ibidi).

### Statistical analysis

All experimental data were plotted and analyzed for statistical significance using the software OriginPro^®^ 2021(OriginLab). Since the data for FA size was right-skewed and not normally distributed, we used the Kruskal –Wallis one-way analysis of variance to compare the different groups. We then used the median FA sizes from each group to plot the FA size trend. The FA data were statistically compared using one-way ANOVA followed by Tukey’s mean comparison, the VE-cadherin data were compared using the unpaired Student’s t-test (two-tailed), and the cell area data was compared using the Mann-Whitney Test.

## General

The authors would like to thank Dr. Anjishnu Sarkar for his assistance in SEM imaging. Cartoons included in the schematic were adapted from Servier Medical Art licensed under a Creative Commons Attribution 3.0 Unported License (smart.servier.com).

## Author contribution

G.P., L.H.R., and D.H.G. conceived the study and designed the experiments. G.P. performed the experiments and data analysis under the supervision of L.H.R., and D.H.G. S.G. helped with schematics and data analysis. J.T. and S.S. formulated the mathematical model. M.N. and G.S. performed the Brillouin imaging. G.P., L.H.R., and D.H.G. wrote the manuscript with input from all the authors.

## Data and materials availability

All data necessary for evaluation of the conclusions in this work are present in the paper and/or the Supplementary Materials. Additional detail is available from authors upon request.

