## Supplementary Material for "Directing multicellular organization by varying the aspect ratio of soft hydrogel microwells"

**Affiliations**

MD 21218, USA.

[14]Department of Chemistry, Johns Hopkins University, Baltimore, MD 21218, USA.

[15]Laboratory for Computational Sensing and Robotics (LCSR), Johns Hopkins University, Baltimore, MD 21218, USA.

**This PDF file includes:**

Supplementary Text

Figs. S1 to S14

Tables S1

### Supplementary Text

#### General model description

To study the mechanism of multicellular organization in microwells, we developed a mathematical model based on force interactions between the cells and substrate. As illustrated in (Fig. 5A and Fig. S13A), the model consists of cells in a hemispherical microwell assumed to be moving upwards. We modeled the cell as a single-line element and assumed that all the forces only acted on each cell's endpoints.

The geometric shape of the microwell with depth $H$ and radius $R$ can be described as:

$$z = H - H\sqrt{1 - \frac{r^2}{R^2}} \qquad (1)$$

where $z$ is the local height which ranges from 0 to $H$ and $r$ is the local radius at $z$, ranging from 0 to $R$. $(r(z = 0) = 0; r(z = H) = R)$. The region outside the microwell is infinitely flat, or: $z = H$ for $r > R$. The angle between the local tangential line (red dotted line in Fig. S12A) and $x$-axis can be described as: $\tan\theta = \frac{dz}{dr} = \frac{Hr}{R^2\sqrt{1-\frac{r^2}{R^2}}}$ for $r < R$; and $\tan\theta = 0$ for $r \geq R$, or $z = H$

By considering cytoskeletal tension ($F_k$), which regulates the cell shape and size, cell-substrate interaction which is the frictional force ($F_\eta$), and cell-cell interaction ($F_w$), we can write the force-balance equation at $i$ th node of $j$ th cell, $\mathbf{x}_{j,i}$, as:

$$\eta\dot{\mathbf{x}}_{j,i} = -\frac{d}{d\mathbf{x}_{j,i}}\left[\frac{k}{2d_0}\left(d_j - d_0\right)^2\right] + A\mathbf{f}_{j,i} + F_0\sum_{m,k}\mathbf{w}\left(\mathbf{x}_{j,i}, \mathbf{x}_{m,k}\right) \qquad (2)$$

where each cell is described by two nodes ($i = 1, 2$). We describe the intracellular cytoskeletal tension using spring potential, with the stiffness of $k$. The equilibrium length of the cell is $d_0$, and the current length of the cell $j$ is $d_j = \left|\mathrm{x}_{j,1} - \mathrm{x}_{j,2}\right|$. The cell constantly generates active protrusion and contraction forces, $\mathrm{f}_{j,i}$. This force, scaled by a constant $A$, follows the random Gaussian distribution with zero mean and is inversely proportional to the cell length, $d_j$: $\mathrm{f}_{j,i} = \frac{E(t)}{d_j}$, $< E(t_1)E(t_2) > = \sigma^2\delta(t_1 - t_2)$ where $\sigma$ is the standard deviation of the Gaussian distribution. This force is along the local tangential direction of the substrate. Interaction between the $i$ th node of the $j$ *th* cell and the $k$ th node of the $m$ th cell is $\mathrm{w}\left(\mathrm{x}_{j,i}, \mathrm{x}_{m,k}\right)$, and follows the van der Waals relation. This interaction becomes repulsive when the distance between the two nodes, $\left|\mathrm{x}_{j,i} - \mathrm{x}_{m,k}\right|$, is less than $s_0$ (see Table S1), attractive if the distance is greater than $s_0$ and fades to zero when $\left|\mathrm{x}_{j,i} - \mathrm{x}_{m,k}\right| \to \infty$. To circumvent the singularity caused by the nature of van der Waals relation when $\left|\mathrm{x}_{j,i} - \mathrm{x}_{m,k}\right| \to 0$, we fitted a continuous stepwise function to the van der Waals equation, so that $\mathrm{w}\left(\mathrm{x}_{j,i}, \mathrm{x}_{m,k}\right)$ goes from -1 to 1 (Fig. S13B). $\mathrm{w}\left(\mathrm{x}_{j,i}, \mathrm{x}_{m,k}\right)$ is scaled by a constant $F_0$, and is along the linear direction between these two nodes (green dotted line in Fig. S13A). $F_0$ represents cell-cell interaction strength, which is a function of VE-cadherin or other intracellular protein activities. We assume the cell-cell interaction is only significant between the top node of cell 1, $\mathrm{x}_{1,2}$ and the bottom node of cell 2, $\mathrm{x}_{2,1}$ since the distances between any other two nodes are much further than $s_0$. $\eta$ is the frictional coefficient between the cell and substrate. The right-hand side of the force balance equation describes the residual force resulting from tension, random protrusions and contractions, and cell-cell interaction, and the left-hand side describes the sliding frictional force between the cell and the substrate. This force balance equation indicates the unbalanced force at each node compensated by the local sliding motion along the microwell.

**Numerical Results**

Since the cells can only move along the tangential direction, we can simplify the system by considering only the tangential direction of equation (2) for each cell node:

$$\begin{aligned}
\dot{x_{1,1}} &= \frac{k}{d_0\eta}(d_1 - d_0)\cos\left(\phi_1 - \theta(x_{1,1})\right) + \frac{A}{d_1\eta}E_{1,1}(t) \\
\dot{x_{1,2}} &= -\frac{k}{d_0\eta}(d_1 - d_0)\cos\left(\phi_1 - \theta(x_{1,2})\right) + \frac{A}{d_1\eta}E_{1,2}(t) - \frac{F_0}{\eta}w_{1,2\to2,1}\cos\left(\beta - \theta(x_{1,2})\right) \\
\dot{x_{2,1}} &= \frac{k}{d_0\eta}(d_2 - d_0)\cos\left(\phi_2 - \theta(x_{2,1})\right) + \frac{A}{d_2\eta}E_{2,1}(t) + \frac{F_0}{\eta}w_{1,2\to2,1}\cos\left(\beta - \theta(x_{2,1})\right) \\
\dot{x_{2,2}} &= -\frac{k}{d_0\eta}(d_2 - d_0)\cos\left(\phi_2 - \theta(x_{2,2})\right) + \frac{A}{d_2\eta}E_{2,2}(t)
\end{aligned} \tag{3}$$

in which $\phi_j$ denotes the angle between $j$ th cell $(\mathrm{x}_{j,2} - \mathrm{x}_{j,1})$ and the $x$-axis. $\phi_j - \theta(x_{j,i})$ denotes the orientation of the cells relative to the microwell's tangential direction at the $i$ th node of the $j$ th cell: $(x_{j,i})$. $\beta$ is the angle between the line connecting the two cell nodes $(\mathrm{x}_{2,1} - \mathrm{x}_{1,2})$ and the $x$-axis. We used a MATLAB package to independently generate random $E_{1,1}, E_{1,2}, E_{2,1}$ $E_{2,2}$ from the normal distribution with mean equals to zero and standard deviation of pre-defined $\sigma$ at each time step.

We computed Eq. 3 for 1,000 timesteps with a time increment of $dt$ = 0.01 and ran the entire simulation over 200 times to statistically quantify the steady-state multicellular organization in the microwell. We determined whether the cells were located edge or to the center by defining a height ratio: $\frac{z_0}{H}$, in which $z_0$ is the average $z$ coordinate of the cells' center point after each of the 1,000-time iteration for each simulation. If $\frac{z_0}{H}$ is close to 1, the cells are more likely to be at the microwell edge, and if $\frac{z_0}{H}$ is closer to zero, the cells are more likely to be towards the center. We define cells to be organized on the edge if $\frac{z_0}{H} \geq 0.8$, and to the center if $\frac{z_0}{H} \leq 0.2$. We then quantified the ratio between the probability of cells staying on the microwell's edge and the probability of the cells staying towards the center, $N_{Edge}/N_{center}$ based on all 200 simulations. We define $N_{Edge}/N_{center}$ as DR in the main text.

The cells are more likely to stay at the microwell center in the shallower microwells but relocate to the edges of deeper microwells. With the correct set of parameters, our simulation results can predict cellular organization as observed experimentally (Fig. 5C). We could also predict the sensitivity of multicellular organization to geometry in response to cytoskeletal contractility (or tension) influenced by substrate stiffness and cell-cell interactions using this force balance model (Fig. 5C and D), and these predictions are consistent with experimental findings (Fig S7, S8, and S10).

The observed spatial localization can be explained by the force-balance feature of the system, caused due to the geometry of the microwell. Cells rely on cytoskeletal activities and forces to move and balance extracellular forces, including cell-cell interactions and cell-substrate friction. Since the cytoskeletal tension is not precisely along the microwell's local tangential direction, the magnitude of tangential components of cytoskeletal tension depends on the cell's orientation relative to the microwell's tangential direction. The unbalanced part of the force drives the cell movement along the microwell. Therefore, the cell is less likely to move when the cytoskeletal tension is more likely to balance the extracellular forces. This balance occurs at microwell positions where $|\phi_j - \theta_{j,i}| \sim 0$. According to the geometric calculation, $|\phi_j - \theta_{j,i}|$ is close to zero around the center of shallower microwells or along the edge of deeper microwells (Fig. S13C). Thus, when the cell-cell interactions are increased, higher tangential intracellular cell tension components are required to balance the increased interaction, which is achievable on the microwell edge. Similarly, when the cell contractility ($k$) is decreased, due to a decrease in substrate stiffness, the cell again moves to the edge where a higher tangential component of tension is achievable to balance the other forces of the system. Further, to quantify the unbalanced force which drives the cell movement up and down, we calculated the mean vertical velocity of the cells when they are located at the center and top edge of the microwell, respectively, and averaged them over all the nodes, $v_z$. The vertical velocity indicates the vertical component of the frictional force, which drives the vertical movement of the cells. As shown in (Fig. S13D), when cells are seeded in deeper microwells, higher vertical velocity towards the center and lower vertical velocity towards the top edge is predicted compared to cells seeded in the shallower microwell, indicating a higher probability of cells moving towards the top edge in deeper microwells ($\varepsilon < 25$), and towards the center in shallower microwell ($\varepsilon > 25$).

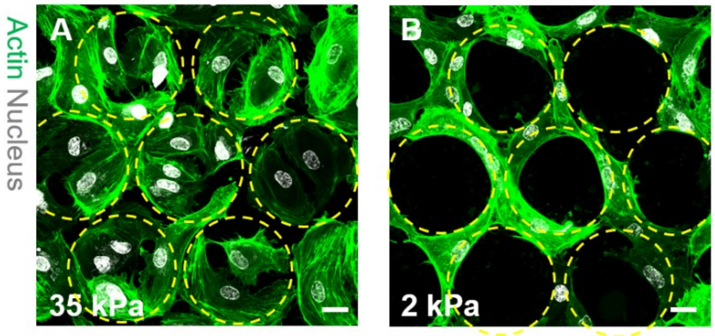


**Fig. S1. Multicellular self-organization similar to that on gelatin microwells was also observed in polyacrylamide hydrogel microwells.** (**A** and **B**) Confocal images showing top view of HUVEC cells labelled for actin (green) and nucleus (gray) on stiff 35 kPa and soft 2 kPa polyacrylamide hydrogels 24 h after seeding. Scale bar = 25 µm.

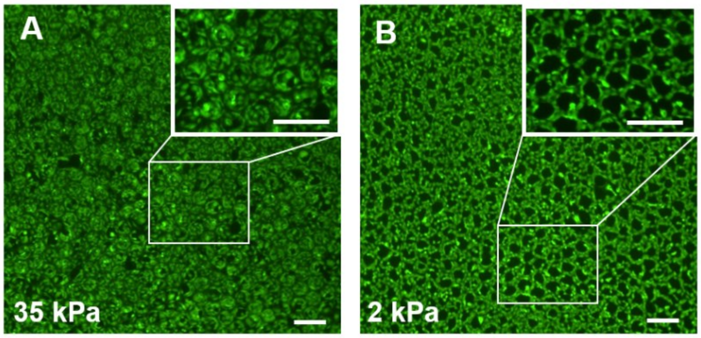


**Fig. S2. Cell self-organization over large areas indicating that the phenomenon and microwell hydrogels are spatially uniform.** (**A** and **B**) Large area (1mmx1mm) fluorescence microscopy images of cells stained with Calcein AM in stiff 35 kPa (A), and soft 2 kPa (B) gelatin microwells 24 h after seeding. Scale bar = 200 µm

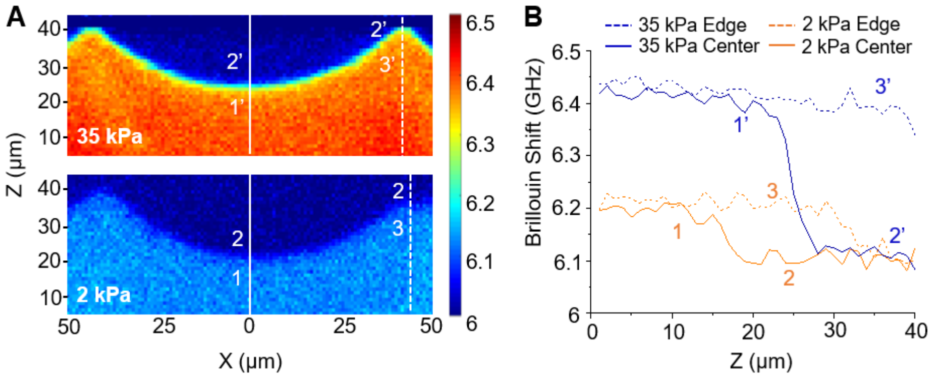


**Fig. S3. Brillouin images indicating homogenous mechanical properties along the z axis.** (**A**) Representative Brillouin images depicting the microwell cross-section for stiff 35 kPa and soft 2 kPa microwells. (**B**) Representative plot depicting the Brillouin frequency shift along the z direction at two positions; microwell center as indicated by the solid line and the microwell edge indicated by the dashed line. Point 1',3' (35 kPa) and 1,3 (2 kPa) are inside the hydrogel whereas points 2' (35 kPa) and 2 (2 kPa) are in PBS. The transition from higher constant value of Brillouin shift to 6.1 GHz (Brillouin shift of the surrounding PBS) corresponds to 2 µm, which is the axial resolution of the objective we used to measure Brillouin shift.

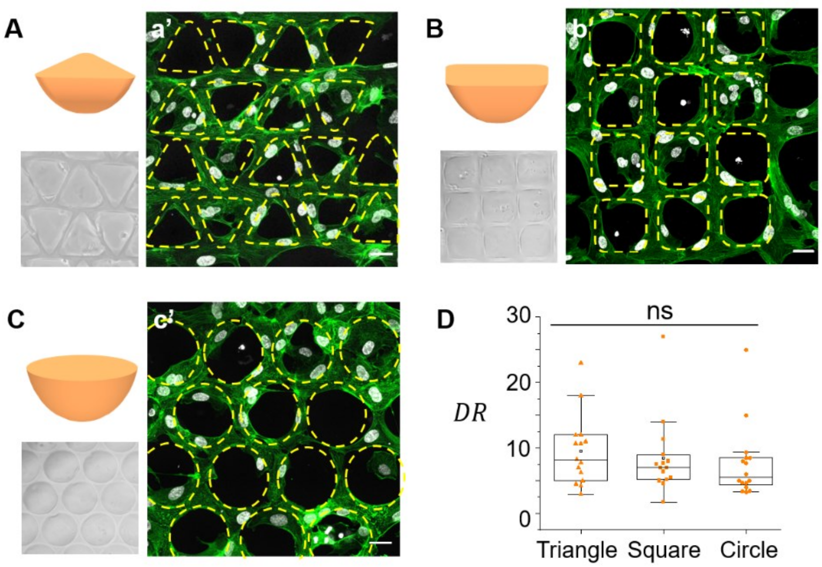


**Fig. S4. Cell organization on soft hydrogels is independent of the microwell shape**. (**A, B,** and **C**) Schematic and optical images of microwells with triangular, square, and circular perimeter. (**A', B',** and **C'**) Confocal images showing top view of cells in microwells corresponding to (A, B, and C). Scale bar = 25 µm. (**D**) Box plot depicting the distribution of cells (DR) in microwells of different shapes.

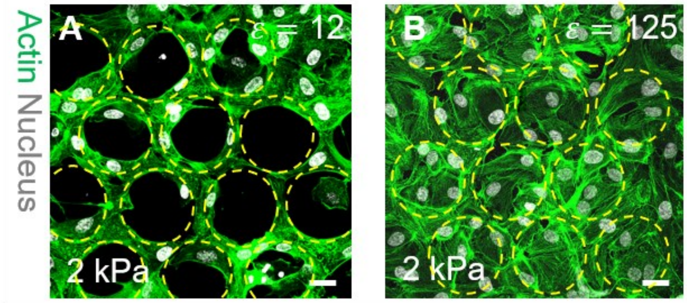


**Fig. S5. Multicellular organization in 2 kPa microwells with two different microwell aspect ratios are dramatically different.** (**A** and **B**) Confocal images showing top view of cells stained for actin (green) and nucleus (gray) in soft microwells with depth 20 µm, $\varepsilon$ =12 (A), and 2 µm, $\varepsilon$ =125 (B) 24 h after seeding. Scale bar = 25 µm. Yellow dashed circles depict the microwell location.

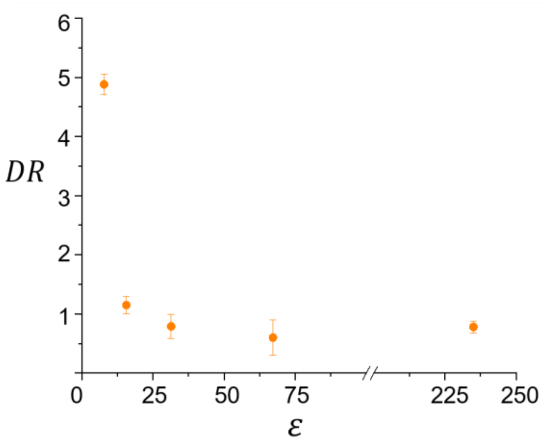


**Fig. S6. Multicellular organization in larger microwells follows a similar trend as in 250 µm perimeter microwells.** Plot depicting extent of self-organization in soft 2 kPa microwells with a perimeter of 470 µm as the aspect ratio ($\varepsilon$) increases (depth decreases). Data presented are quantified from 100 cells for each aspect ratio and repeated three times. Error bars indicate standard deviation between each experiment.

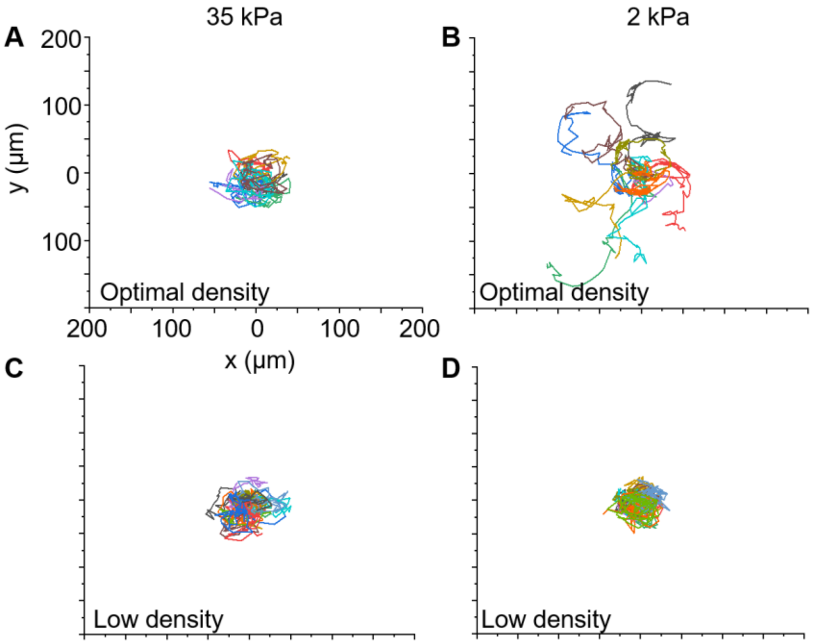


**Fig. S7. Migration of cells with and without neighbors in the microwell.** (**A** to **D**) Cell migration tracks in stiff 35 kPa and soft 2 kPa microwells at optimal cell density (A and B) and low density (C, D). At optimal seeding density, each microwell has 3-5 cells allowing them to make cell-cell adhesions whereas at low density each microwell has one cell.

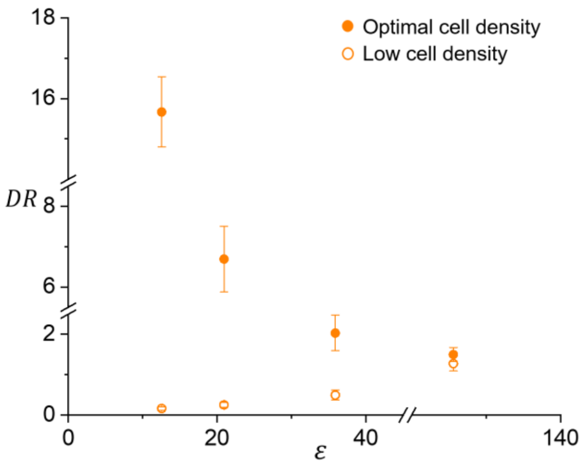


**Fig. S8. Cells are predominantly found at the center of the soft 2 kPa microwells at low cell density.** Plot depicting cell distribution at optimal (solid circles) and low (open circles) cell density in soft 2 kPa microwells in relation to the microwell aspect ratio ($\varepsilon$). Data presented are quantified from 100 cells for each aspect ratio and repeated three times. Error bars indicate standard deviation between each experiment.

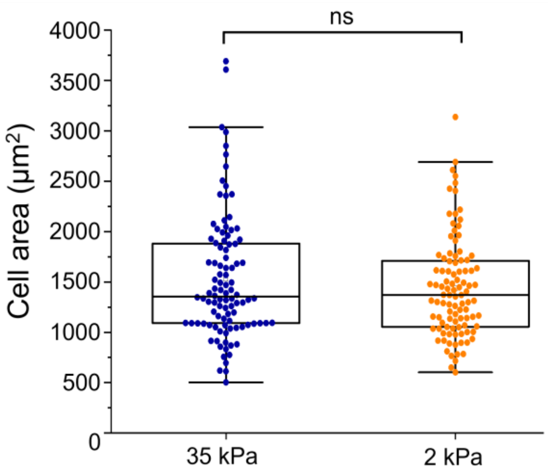


**Fig. S9: Area of cells in 35 kPa and 2 kPa microwells is similar**. Plot depicting the projected area of cells in 35 kPa and 2 kPa microwells measured by projecting the z-stack on a plane. Data presented are quantified from 100 cells selected from three independent experiments. *P* value >0.05 as calculated using Mann-Whitney test.

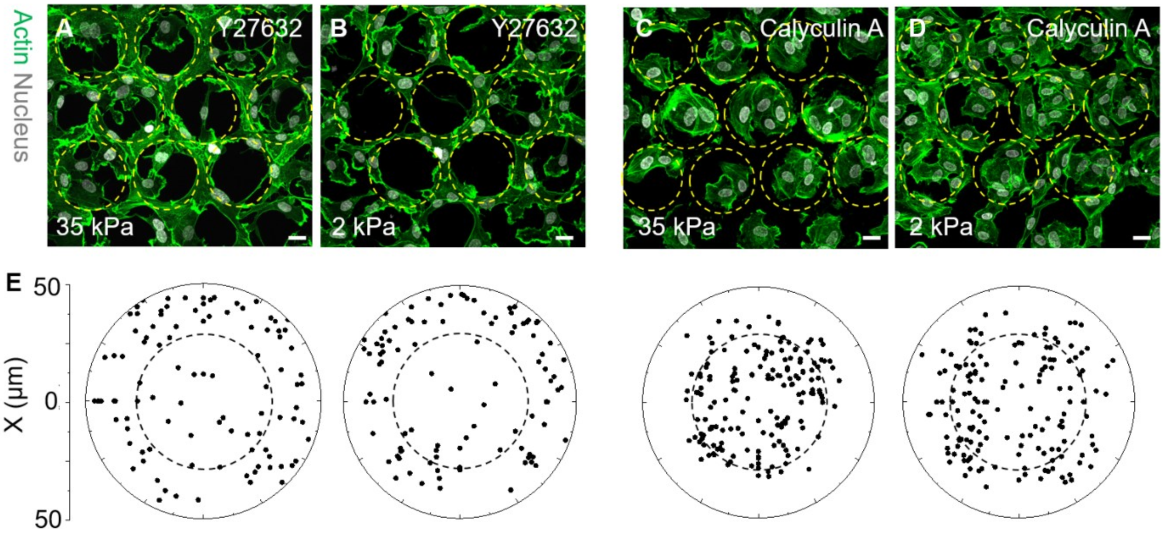


**Fig. S10. Lower cell cytoskeleton contractility enhances self-organization. (A** and **B)** Confocal images showing top view of Y27632 treated cells stained for actin (green) and nucleus (gray) in stiff 35 kPa (A), and soft 2 kPa microwells (B). (**C** and **D**) Confocal images showing top view of Calyculin A treated cells actin (green) and nucleus (gray) in stiff 35 kPa (C), and soft 2 kPa microwells (D). Scale bar = 25 μm. Yellow dashed circles depict the microwell location. (**E**) Polar plot depicting the distribution of cells in the microwell corresponding to conditions in (A, B, C, and D). Data presented are for 100 cells in each condition pooled from three independent experiments.

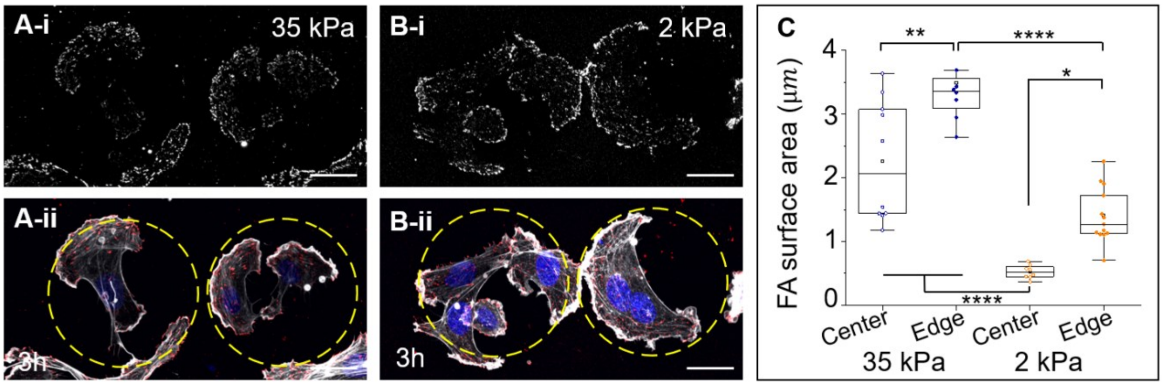


**Fig. S11. FA distribution in microwells 3 h after seeding.** (**A** and **B**) Confocal microscopy images showing the top view of cells immunostained for paxillin (gray) 3 h after seeding in stiff 35 kPa (A-i) and soft 2 kPa (B-i) microwells and the corresponding composite top view of cells stained for paxillin (red), actin (gray) and nucleus (blue) (A, B-ii). Yellow dotted circles indicate the microwell position. Scale bar: 25 µm. (**C**) Box plot depicting paxillin containing FA surface area at the center and edge of stiff 35 kPa and soft 2 kPa microwells. Data presented are for n ≥8 cells pooled from 3 independent experiments. ****$P$ value <0.0001 and **P <0.01 calculated by one-way ANOVA followed by Tukey's multiple comparison.

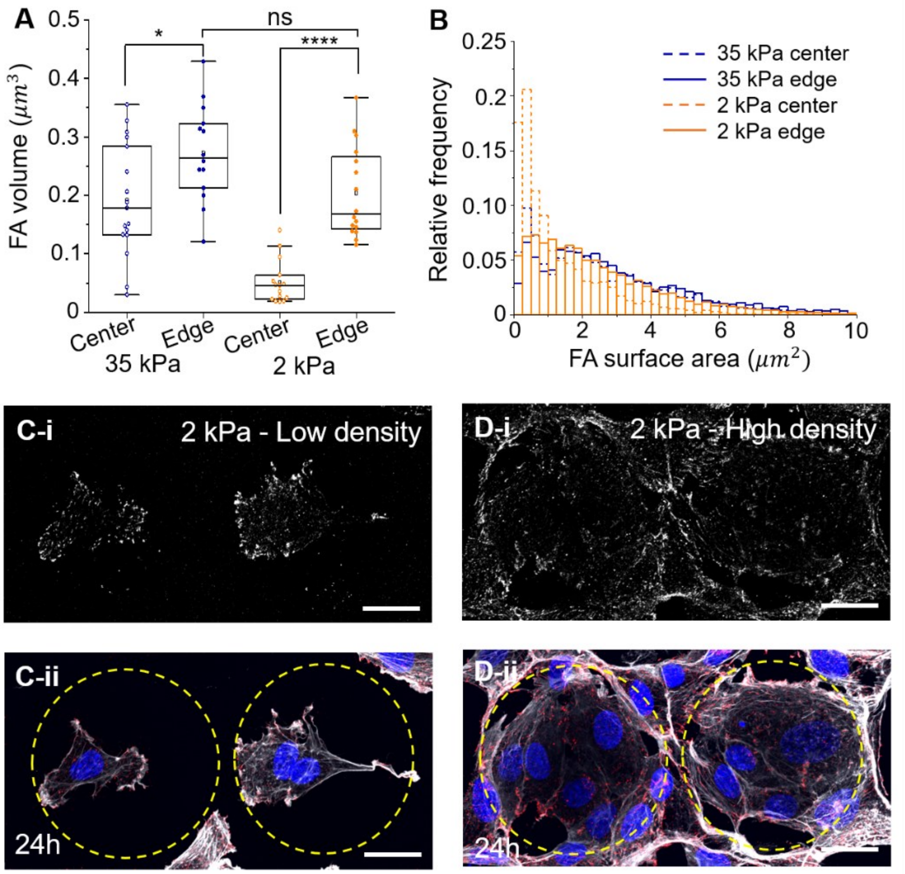


**Fig. S12. FA distribution in microwells 24 h after seeding.** (A) Box plot depicting FA volume at the center and edge of stiff 35 kPa and soft 2 kPa microwells. **P* value <0.01 and ****P value < 0.0001 calculated from one-way ANOVA followed by Tukey's multiple comparison. **(B)** Histogram depicting the relative frequency of FA of a particular size**. (C and D)** Confocal microscopy images showing the top view of cells in soft 2 kPa microwells immunostained for paxillin 24 h after seeding at low density (C-i) and high density (D-i) and the corresponding top view of cells stained for paxillin (red), actin (gray), and nucleus (blue) (C, D-ii). Yellow dotted circles indicate the microwell position. Scale bar = 25 μm.

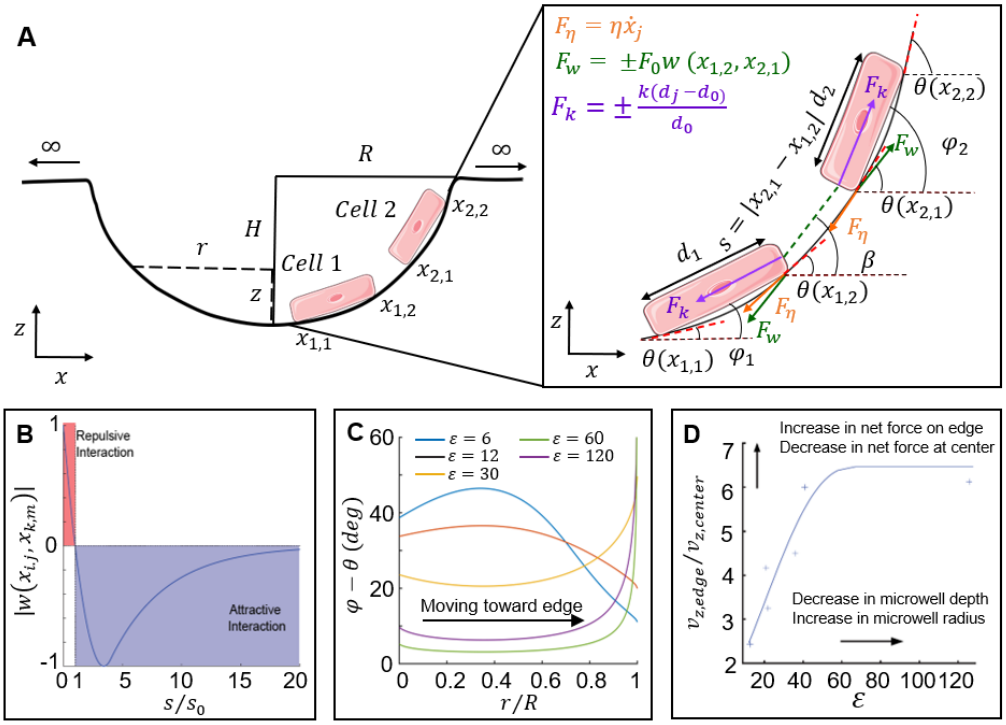


**Fig. S13. Force balance model description. (A)** Schematic depicting the force balance model used to simulate the cell organization. **(B)** Cell-cell attractions along the linear direction modeled using the van der Waals relation. **(C)** Orientation of the cell relative to the microwell tangential direction at different microwell locations for various aspect ratios. **(D)** Plot depicting the vertical frictional force as a function of the microwell aspect ratio ($\varepsilon$).

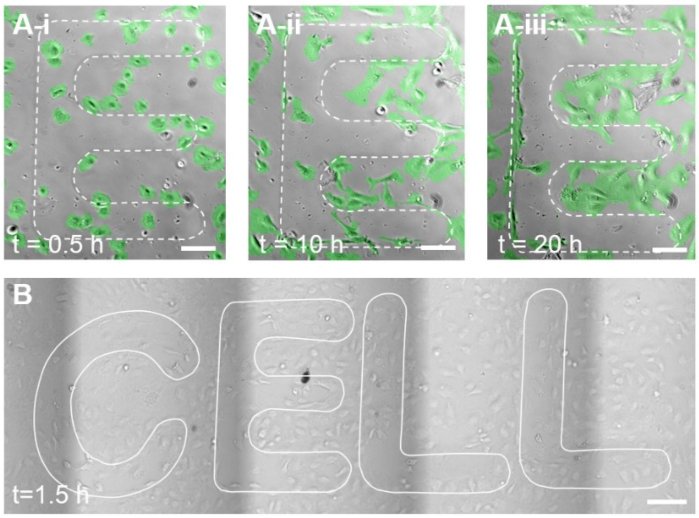


**Fig. S14. Cell organization on soft hydrogels micropatterned with the word CELL. (Ai-iii)** False colored time-lapse sequence of cells moving in the letter E. **(B)** Bright-field image depicting cells on the pattern CELL 1.5 h after cell seeding. Scale bar = 100 µm.

| Parameters | Description | Value/Range |
| --- | --- | --- |
| $\varepsilon = \frac{2\pi R}{H}$ | Ratio between microwell perimeter and depth | $2\pi \times (1 \sim 20)$ |
| $\frac{d_0}{R}$ | Scaled equilibrium length of the cell | 0.5 |
| $\frac{s_0}{d_0}$ | Scaled equilibrium distance of van der Waals potential | 0.1 |
| $\sigma$ | Standard deviation of random protrusions/contractions distribution | 1 |
| $\frac{kdt}{\eta d_0}$ | Scaled spring constant | 0.6~0.8 |
| $\frac{F_0 dt}{\eta d_0}$ | Scaled cell-cell interaction strength | 1.5 ~ 3 |
| $\frac{Adt}{\eta d_0}$ | Scaled random protrusions/contractions | 1 |
| $dt\ (s)$ | Time increment | 0.01 |

**Table 1.** Description of parameters used in the force-balance model.
